# A network model of the modulation of gamma oscillations by NMDA receptors in cerebral cortex

**DOI:** 10.1101/2021.12.21.473671

**Authors:** Eduarda Susin, Alain Destexhe

**Author notes:** ES and AD designed the study and the model, ES performed simulations and analysis, ES and AD discussed the results and wrote the paper, AD supervised the study. Authors declare no competing interest.

## Abstract

Psychotic drugs such as ketamine induce symptoms close to schizophrenia, and stimulate the production of gamma oscillations, as also seen in patients, but the underlying mechanisms are still unclear. Here, we have used computational models of cortical networks generating gamma oscillations, and have integrated the action of drugs such as ketamine to partially block n-methyl-d-Aspartate (NMDA) receptors. The model can reproduce the paradoxical increase of gamma oscillations by NMDA-receptor antagonists, assuming that antagonists affect NMDA receptors with higher affinity on inhibitory interneurons. We next used the model to compare the responsiveness of the network to external stimuli, and found that when NMDA channnels are diminished, an increase of gamma power is observed altogether with an increase of network responsiveness. However, this responsiveness increase applies not only to gamma states, but was also present in asynchronous states with no apparent gamma. We conclude that NMDA antagonists induce an increased excitability state, which may or may not produce gamma oscillations, but the response to external inputs is exacerbated, which may explain phenomena such as altered perception or hallucinations.

**Significance Statement:** n-methyl-d-Aspartate (NMDA) synaptic receptors mediate excitatory interactions using the neurotransmitter glutamate. NMDA receptors have been implicated in psychosis such as schizophrenia and are also targeted by hallucinogenic drugs like Ketamine. However, the exact mechanisms of action are sill unclear. Furthermore, Ketamine paradoxially leads to and excited state, while it is a blocker of NMDA receptors, therefore in principle diminishing excitation. Here, we use models of cortical networks generating gamma oscillations, and show that this model can explain the paradoxical exciting effect of Ketamine if one assumes a higher affinity on NMDA receptors of inhibitory interneurons. The simulated Ketamine effect reproduces known symptoms of psychosis such as increased gamma oscillations and exacerbated responses to external inputs, compatible with hallucinations.

---

Schizophrenia is a mental disorder characterized by three classes of symptoms: positive symptoms (such as delusions, hallucinations and disordered thoughts or speech), negative symptoms (comprehending poverty of speech and deficits of normal emotional response), and cognitive deficits (1–3). Several abnormalities have been identified in schizophrenic patients, including important differences in neurotransmitters systems, anatomical deficits and abnormal neural rhythms (4, 5).

Gamma oscillations (30-90 Hz) in early-course schizophrenia patients are commonly reported to present increased power and/or phase synchronization (6–8). In parallel, positive correlation between psychotic symptoms and the gamma power have been identified in schizophrenic patients, in which higher gamma-band activity corresponded to increased symptom load (9–12). These findings indicate that hallucinations and delusions could be related to an excess of oscillatory synchronization in the gamma band.

NMDA receptor (NMDAR) antagonists, commonly used in sub-anesthetic doses as animal and human models to study Schizophrenia (13), induce a psychotic state that resembles all three classes of symptoms of the disease (14–16). Furthermore, NMDAR antagonists also increase gamma power amplitude, both in human and in animal models (17–24).

In this study we investigate by means of computational models how NMDAR antagonists, such as ketamine, affect the dynamics of neural networks and how the generated boosting of gamma activity affects the network response, providing an interpretation for the observed correlation between gamma power and psychotic episodes.

## Results

### Modeling the effect of NMDA receptor antagonists

Several studies using sub-anesthetic doses of NMDAR antagonists have reported to produce neural excitation (25–30). Since NMDAR mediate excitatory synaptic transmission, this behavior is paradoxical. Several hypothesis have been proposed to explain this apparent paradox (3). One of the possible explanations is that NMDAR antagonists in sub-anesthetic doses act preferentially on inhibitory neurons, increasing network activity indirectly by means of desinhibition. Even though some contrasting results have been reported (31), this interpretation has been supported experimentally by several works (32–35). Network excitability have also been reported to increase in schizophrenic patients (36, 37), and its increase in sensory and association cortex have been correlated with hallucinations (38, 39).

Another important effect of NMDAR antagonists in subanesthetic doses is the increase of gamma-band activity. These observations were reported in human (17–19), monkey (24) and rats (20–23), both during cognitive tasks or free movement.

The network model developed in the present work (see Methods) is able to reproduce both of these features (increase of network excitability and increase of gamma power). Fig. 1 depicts the network behavior with respect to the to different NMDA synaptic strengths, *Q^NMDA^*, in excitatory Regular Spiking (RS) and in inhibitory Fast Spiking (FS) cells. We mimic the block of NMDA channels due to the action of NMDAR antagonists by decreasing *Q^NMDA^* in RS and FS cells according to Fig. 1A (see Methods). Points of higher synaptic strengths are associated with healthy conditions, while points with lower synaptic strengths are associated to pathological conditions supposedly similar to the schizophrenic brain. The network dynamics for two sets of NMDA synaptic strengths are shown in Fig. 1B and Fig. 1C by means of a Raster Plot. As the synaptic strengths of NMDA channels decreased (higher concentration of NMDAR antagonists), the firing rate of excitatory RS cells increased while the firing rate of inhibitory FS cells decreased (Fig. 1D). This was paralleled with an increase of gamma power of the population activity (Fig. 1E and F; see Methods).

**Fig. 1.**
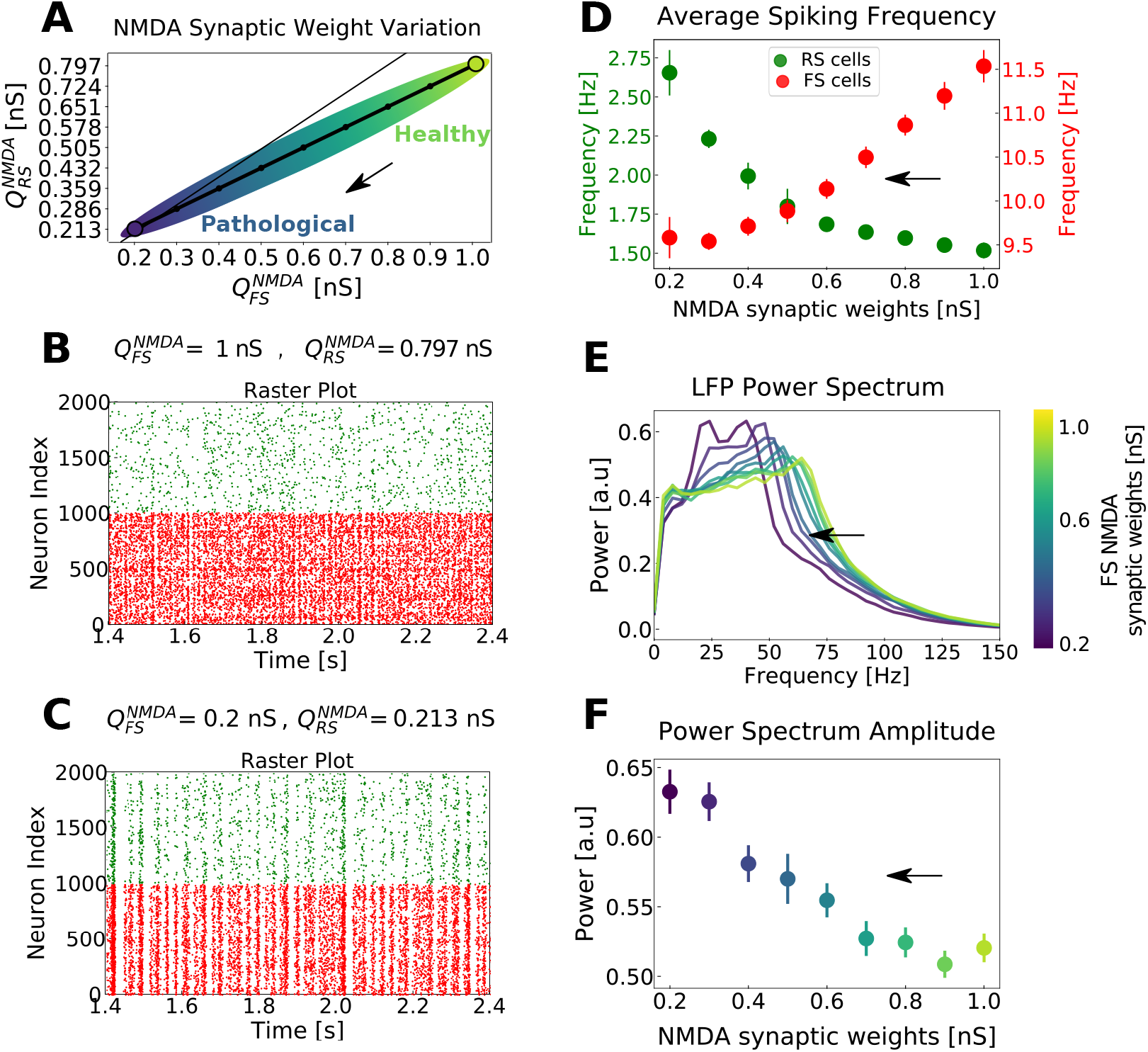
Network dynamics with respect to different levels of NMDA channels block in the network. A) Possible trajectory in the parameter space of 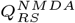 vs. 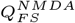, mimicking the action of NMDA receptor (NMDAR) antagonists (the higher the intensity of the NMDAR antagonists, the smaller the NMDA synaptic strengths). The thin line indicates the identity for reference. The arrow indicates the sens of action of NMDAR antagonists. Points of higher synaptic strengths are associated with healthy conditions, while points with lower synaptic strengths are associated to pathological conditions supposedly similar to the schizophrenic brain. B) and C) Raster plots indicating the activity of only 1000 cells of each type (FS in red and RS in green), for two parameter sets. B: 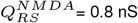 and 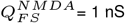, and C: 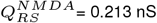 and 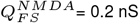. D) Average firing rate of RS (green) and FS cells (red) with respect to the trajectory in parameter space depicted in A. Only the values of 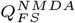 are indicated in the x axis. Standard errors of the mean (SEM) are indicated as error bars. E) Average normalized Power Spectrum of the network LFP for different NMDA synaptic strength. Like in D, the synaptic strengths follow the curve indicated in A, but only the values of 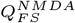 are indicated in the color scheme. Notice the shift of the Power Spectrum peak toward smaller frequencies with the increase of NMDA channel block. F) Power Spectrum peak amplitude with respect to the levels of NMDA channels block (following the synaptic strengths indicated in A). The color scheme (presented for better visualization) are the same as in E. Standard errors of the mean (SEM) are indicated as error bars. Results expressed in D, E and F are the outcome of 50 simulations average. The arrows indicate the sense of the behavior according to amount of block of NMDA channels.

### Network Responsiveness during gamma rhythms with different levels of NMDAR block

We investigated how the decrease of NMDA synaptic strength changed the network dynamics and its capacity to respond to external stimulus.

While network *excitability* is related to an overall increase of spiking activity, network *responsiveness* relates to the network capacity to react to a certain stimulus, producing additional spikes then the ones generated by spontaneous activity. These two dynamical measurements (excitability and responsiveness) are not always congruent, meaning that it is possible to observe an increase in excitability but a concomitant decrease in responsiveness (40).

Network responsiveness was defined as the difference between the total number of spikes generated by the whole network in the presence and in the absence of the stimulus (see Eq 6). We measured network responsiveness at different levels of NMDAR block for different stimulus amplitudes (Fig. 2). The stimulus consisted of a variation in time of the external Poissonian drive, in a Gaussian manner (see Methods).

**Fig. 2.**
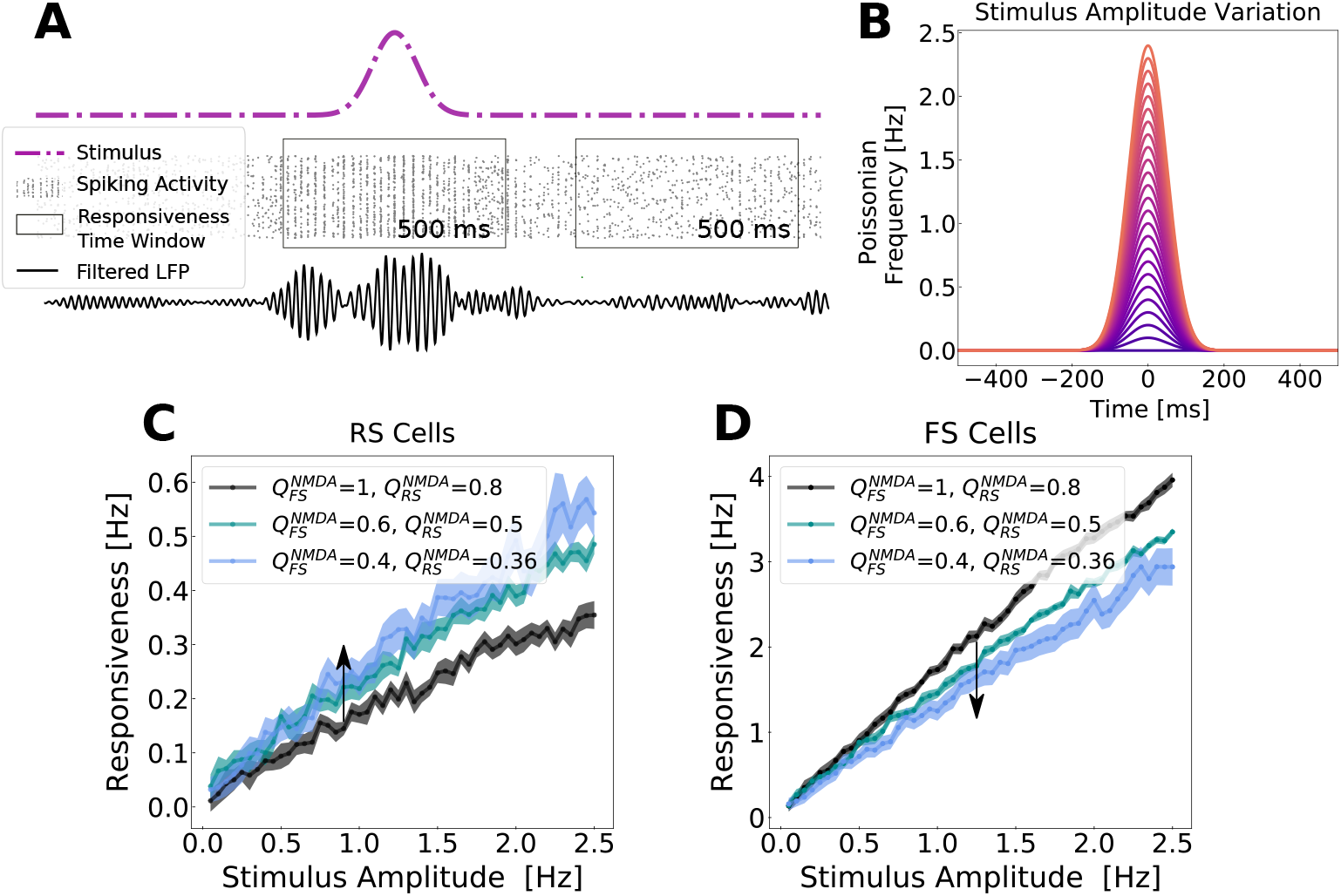
Network responsiveness to broad Gaussian inputs in different levels of NMDA channel blocked during gamma rhythms. A) Responsiveness protocol scheme. The total number of spikes generated by the network were measured during an external stimulus and in its absence in a time window of 500 ms. The stimulus consisted of a Gaussian fluctuation in the firing rate of the external noise input. Responsiveness was calculated according to Equation 6. B) Gaussian input amplitude variation. The Gaussian amplitude varied from 0.05 Hz to 2.5 Hz (step of 0.05 Hz), always keeping the same standard deviation of 50 ms. C and D depict respectively the responsiveness of RS (C) and FS (D) neurons for different Gaussian amplitudes in different levels of NMDAR block, when the network was displaying gamma activity. The color-scheme indicates the synaptic weights of NMDA synapses (*Q^NMDA^*) in RS and FS cells. The arrow indicates the sense of the simulated action of NMDA antagonist (decreasing synaptic strength). Every point corresponds to the average responsiveness measured in 15 simulations. Standard error of the mean are indicated by the shaded region around each curve.

Network responsiveness in RS cells increased with the increased level of NMDAR block, while the responsiveness of FS neurons decreased. In this case, both, network excitability and network responsiveness, behave in the same direction.

The increase of network responsiveness can be understood from Fig. 3. The NMDA receptors block depolarizes RS cells, while FS neurons are overall hyperpolarized. For weak levels of NMDA receptors block, no or weak depolarization is observed in FS cells, while for strong levels of NMDA block a significant hyperpolarization is observed.

**Fig. 3.**
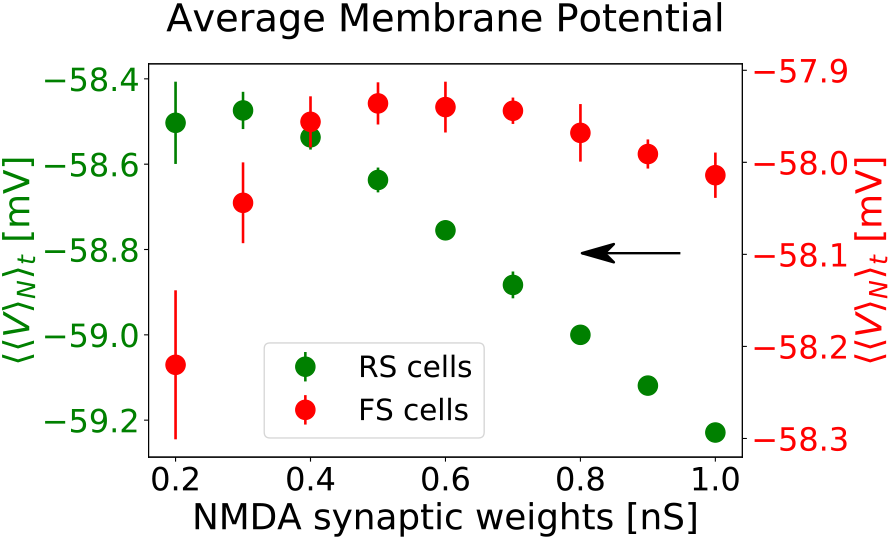
Membrane potential polarization as a function of NMDA receptor block. The average membrane potential of RS (green, left y-axis) and FS (red, right y-axis) is expressed as function of NMDA synaptic weights of RS and FS cells. The values of 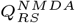 and 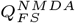 follow the trajectory in the parameter space indicated in Fig. 1A. Only the values of 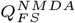 are indicated in the x axis. The average was performed first in between neurons (〈〉*_N_*), obtaining an average curve as a function of time, and subsequently with respect time (〈〉*_t_*). The values plotted correspond to the average of 〈〈*V*〉*_N_*〉*_t_* in between 10 simulations. The error bars indicate the standard error of the mean between these simulations.

### Gamma states vs. AI states

Gamma oscillations (30-90 Hz) are believed to be involved in information processing (41–46), and have been associated to different high-level cognitive functions, such as memory (47–49), perception (50–53), attention (54–57), focused arousal (58) and prediction (59). In parallel, studies with schizophrenic patients have reported a positive correlation between psychotic symptoms and the power of gamma oscillations (9–12).

In contrast, Asynchronous-and-Irregular (AI) states (60) are usually associated to conscious states (61), being observed during awake and aroused states (62). This regime are characterized by irregular and sustained firing with very weak correlations (63–67).

In a previous study (40) we reported that AI states, in comparison to oscillatory states in gamma band, provide the highest responsiveness to external stimuli, indicating that gamma oscillations tend to overall diminish responsiveness. This observation could indicate that gamma rhythms present a *masking effect*, conveying information in its cycles on spike timing at the expense of decreasing the strength of the network response.

In the present study, we compare AI and gamma states at different levels of NMDAR block. Fig. 4 depicts the responsiveness of RS neurons, with respect to different stimulus amplitudes (same protocol as Fig. 2), for different ensembles of NMDA synaptic strengths. In agreement with Fig. 2, parameter sets in which NMDA synaptic strengths are decreased (mimicking the action NMDAR antagonists) correspond to regions of the parameter space with higher responsiveness. For example, 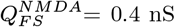 and 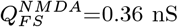 displayed higher responsiveness then the networks in which the NMDA synaptic strengths wrere 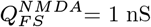 and 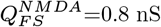. Interestingly, in both conditions, responsiveness in AI states were always superior to the one in gamma. This result was also observed in a similar model in the obscene of NMDA channels (40). This example illustrates a general tendency, which was also observed with other parameter sets.

**Fig. 4.**
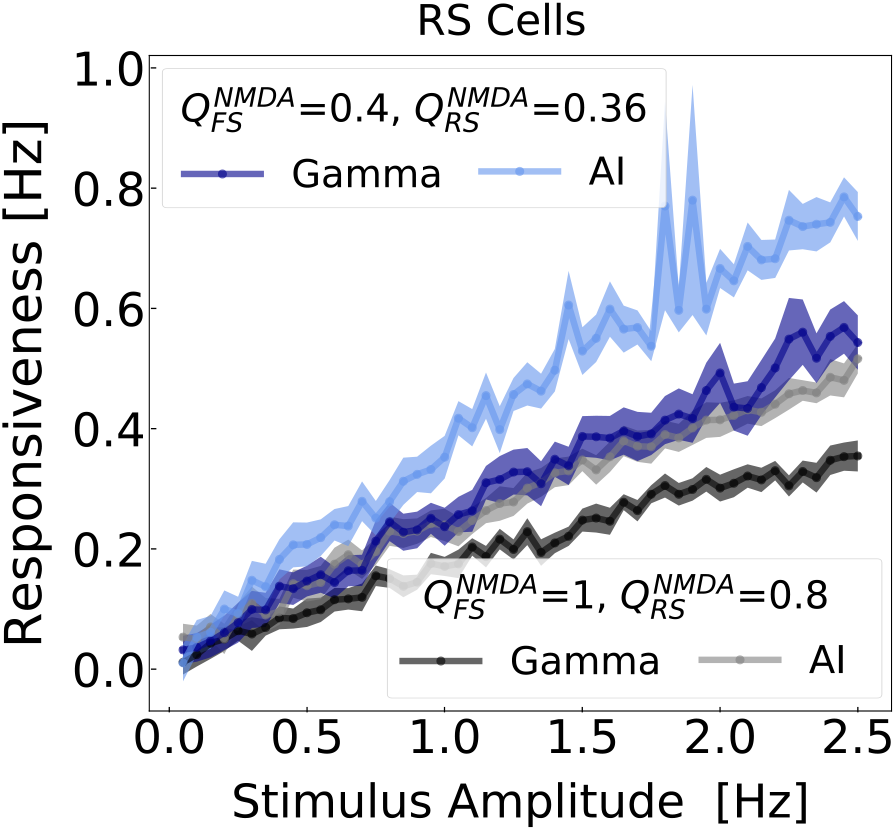
Network responsiveness to broad Gaussian inputs of different amplitudes during gamma and AI states. The responsiveness of RS neurons, due to different Gaussian amplitudes stimuli (same as in the protocol of Fig. 2), was measured in different states AI and gamma for NMDA synaptic parameter sets: 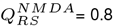 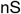 and 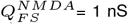 (Gamma: black, AI: gray), and 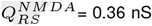 and 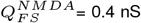 (Gamma: blue, AI: light blue). The Gaussian amplitude varied from 0.05 Hz to 2.5 Hz (step of 0.05 Hz), always keeping the same standard deviation of 50 ms.

### Robustness of the model

As mentioned above, the model was very robust to changes of parameters, and the robustness of the network to produce AI states or gamma oscillations was investigated in detail previously (40). To further characterize its robustness, we represented different aspects of network dynamics as a function of the NMDA synaptic weights on RS and FS cells (Fig. 5). These aspects were the average firing frequency of RS and FS cells, the peak frequency and amplitude in the power spectrum. The paths indicated by white squares and arrow represents a possible trajectory in parameter space following the action of NMDAR antagonists.

**Fig. 5.**
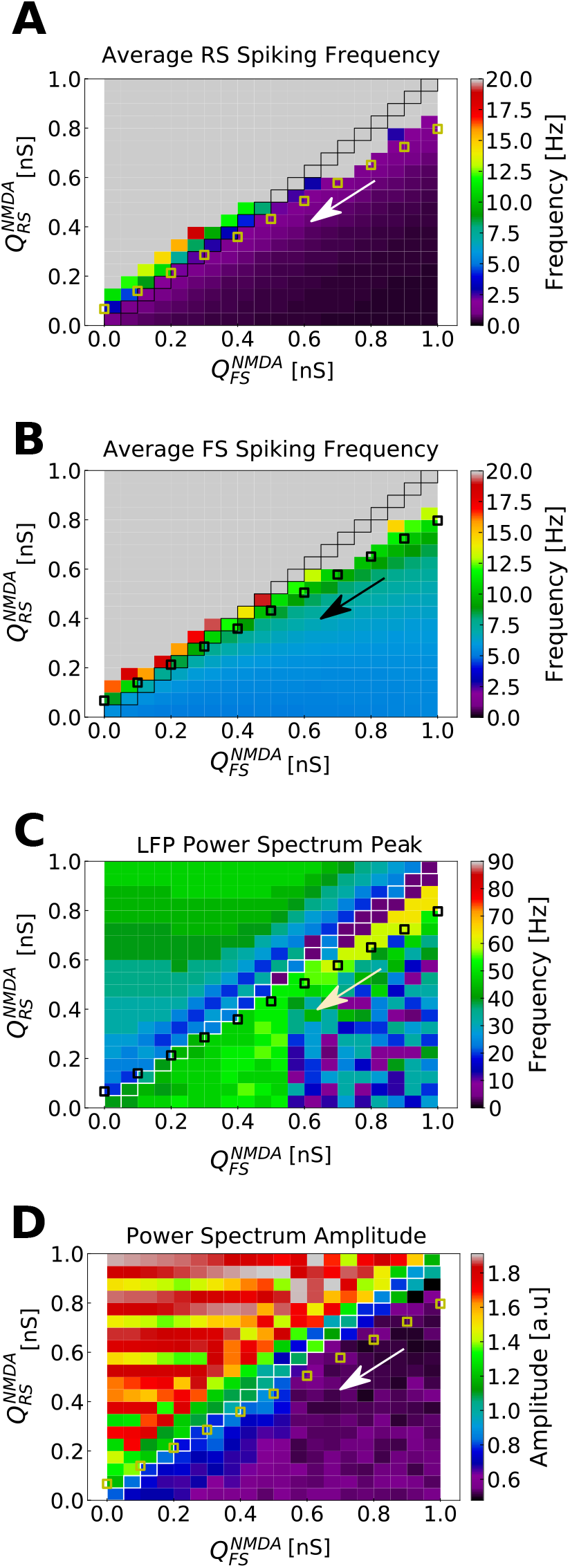
Parameter space of NMDA synaptic weights in RS and FS cells. A) Average spiking rate in RS cells. B) Average spiking rate in FS cells. C) LFP Power Spectrum peak. D) LFP Power Spectrum amplitude. The parameter space of NMDA synaptic weights (*Q^NMDA^*) was explored for RS and FS cells in the developed network model. 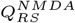 and 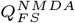 varied from 0 nS to 1 nS in steps of 0.05 nS. Each point in the color maps corresponds to the average of 10 simulations of 5 seconds. Points in which 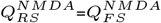 are highlighted. Small squares indicate a possible trajectory in the parameter space (in the direction of the arrow) generated by the action of NMDAR antagonists. This is the same trajectory indicated in Fig. 1A.

The search in parameter space, expressed in Fig. 5, was the basis for the choice of the synaptic strengths of receptors (in RS and FS neurons). With these choice of parameters, the NMDA/AMPA charge ratio in the network is on average higher in RS cells then in FS cells (see Fig. 6), in agreement with experimental measurements in prefrontal cortex of adult mice (31) and rat (68).

**Fig. 6.**
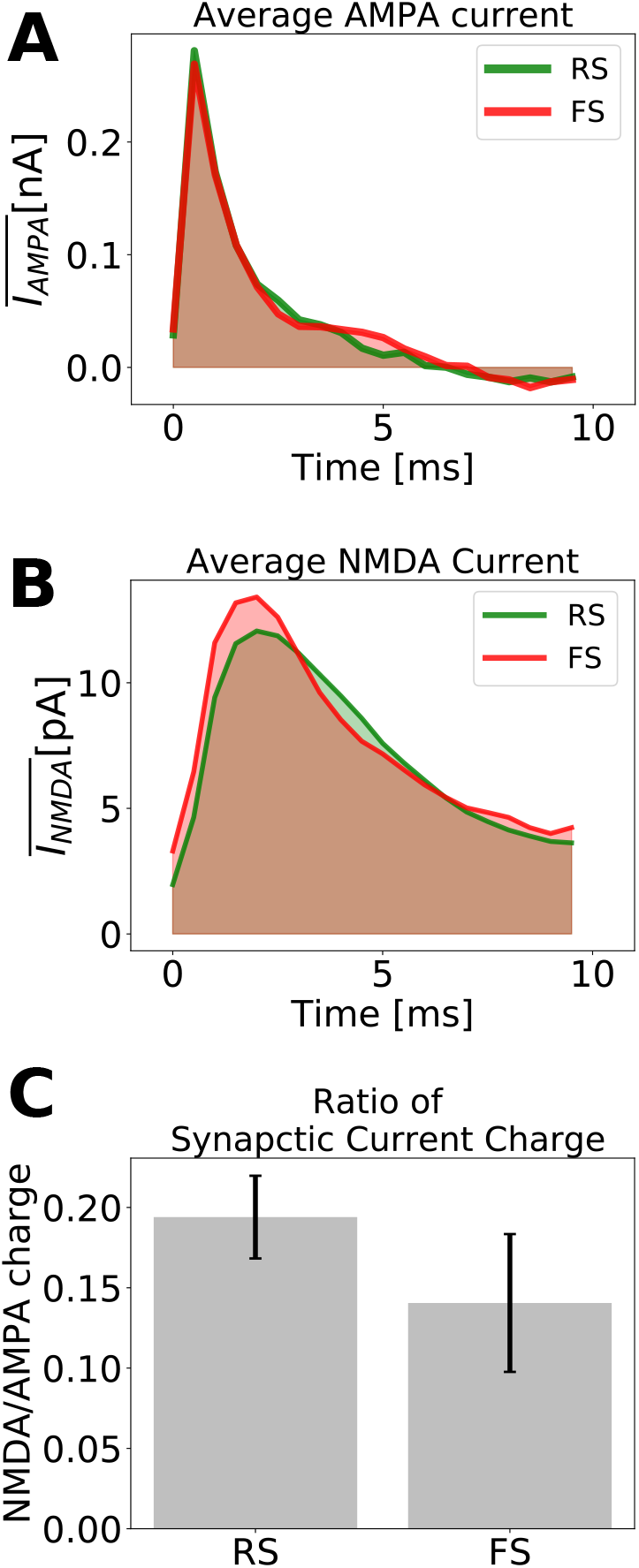
Excitatory synaptic currents. A) Average AMPA current of one randomly picked RS (green) and one randomly picked FS (red) neuron. B) Average NMDA current of one randomly picked RS (green) and one randomly picked FS (red) neuron. C) Ratio of NMDA and AMPA charges for RS and FS cells. The synaptic charge ratio of each neuron was calculated separately. The Bars indicate the mean and the standard deviation among the RS and FS population. The NMDA synaptic strengths in RS and FS cells are 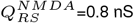 and 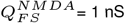 (which, in our model, describes a healthy condition).

## Discussion

In this work, we used computational models to investigate the effect of psychotic drugs such as ketamine in cerebral cortex, and how gamma oscillations relate to these effects. We used a relatively standard model structure to generate gamma oscillations based on two distinct cell types, RS and FS cells (40, 60, 69–75) (although gamma oscillations can also be generated by interneuron networks in some conditions (76, 77); see review in (78)). We integrated NMDA receptors in excitatory connections, treating independently the NMDA input to RS and FS cells. Our findings are (1) NMDA receptors antagonists modulate the rhythms produced by this simple network model. In particular, a boosting of gamma oscillations is obtained assuming that the NMDAR block predominantly affects FS interneurons. (2) The boosted gamma oscillations following partial block of NMDA receptors, was accompanied by an increased responsiveness to external inputs. (3) This increase of responsiveness could also be seen for asynchronous states, with no apparent gamma. We discuss below the implications of these findings.

A first prediction of the model is that it was necessary that the antagonism affects predominantly NMDAR receptors on interneurons. This feature is supported by a number of observations. Intuitively, if the NMDAR block would occur predominantly on excitatory cells, then it is difficult to see how diminishing excitation could augment the activity and excitability of the network. This long-standing question was resolved recently by finding that indeed, NMDAR antagonists primarily affects NMDA receptors on interneurons. It was observed that the application of Ketamine or MK-801 in subanesthetic doses leads to an increased activity of glutamatergic neurons both in cortex (25, 35) and in hippocampus (33), and that this increase of glutamatergic activity is a consequence of the disinhibition of GABAergic neurons (32, 34). In addition, it has also been reported in hippocampus that inhibitory neurons are more sensitive to NMDAR antagonists than glutamatergic neurons (79, 80). Thus, our model completely supports these findings, and could reproduce the increase of gamma power induced by NMDA receptor antagonists. On the other hand, contrasting results also exist. For example, (31) argue that NMDAR have less impact on the activity of inhibitory neurons than on the one of excitatory neurons, since they and other authors observed that NMDAR block depressed large EPSP–spike coupling more strongly in excitatory than in inhibitory neurons (31, 81, 82).

The second finding, which is probably the main finding of our study, is that the network has a marked increased responsiveness under the boosted gamma condition. This increased responsiveness could be tested experimentally either *in vitro*, by testing the response of cortical slices with and without application of NMDAR antagonists, or *in vivo*, by monitoring their response following administration of NMDA antagonists.

The third finding, and prediction, is that the increase of responsiveness is not specific to gamma oscillations, because it was also present for asynchronous states with no apparent gamma. The underlying mechanism is that the antagonism of NMDA receptors produce an overall depolarization of RS cells, and hyperpolarization of FS cells. Consequently, there is an increase of responsiveness of RS cells, with a corresponding decrease for FS cells, as we observed. In this model, the increase of responsiveness is due to the depolarizing effect on RS cells, and are not due to gamma oscillations. Indeed, the highest responsiveness was seen for asynchronous states, also in agreement with a previous modeling study (40).

### Possible implications to understand brain pathologies

Our model exhibits several interesting properties that can be related to pathologies. First, the model provides a possible explanation for the symptoms associated to ketamine and others NMDA receptor antagonists, such as hallucinations. The enhanced responsiveness produced by antagonizing NMDA receptors may explain exacerbated responses to sensory stimuli, which may be related to phenomena such as altered perception or hallucinations. Indeed, it is well documented that ketamine produces hallucinations together with a marked increase of gamma oscillations (83–85).

Besides hallucinations, the model also seems a priori consistent with the previously reported role for FS neurons in schizophrenia. Post-mortem analysis of schizophrenic patient brains have shown a reduced expression of parvalbumin (PV) and GAD67 (1, 86–90). In parallel, genetic ablation of NMDA receptors in PV-positive interneurons in rodents mimics important behavioral (91) and phenotypical features of of the disease (reduction of GAD67 (92), increase of neuronal excitability (92) and increase of spontaneous gamma power (93–95)). These observations support the idea that the hypofunction of NMDA receptors in PV-positive interneurons are specially important in this illness.

However, NMDA receptors are expressed in both GABAergic and glutamatergic neurons (32), and it still remains unclear in which types of cells the NMDA receptor hypofunction would cause schizophrenia (3, 96). Some works reported conflicting results and have questioned the hypothesis that PV-positive Fast Spiking neurons play a role in Schizophrenia (31, 96).

In our model, the effect of NMDAR antagonists is to increase excitability due to disinhibition, consistent with a number of experimental observations (25–30). This increased excitability is accompanied by a gamma power increase, as also found in experiments with ketamine (17–19) or in schizophrenic patients (6–12). The model could reproduce all these experimental observations only assuming a larger decrease of the NMDA synaptic strengths in FS cells than in RS cells (see Fig. 1A). These results support the idea sustained by some authors (97), that PV-positive Fast Spiking inhibitory neurons play a key role in schizophrenia. Another modeling study also stressed the importance of NMDA channels into FS neurons (98). Thus, models support the view that the hypofunction of NMDA receptors on FS cells could explain a number of features typical of schizophrenia, such as anomalous responses and boosted gamma oscillations.

## Materials and Methods

### Neuronal Model

Neural units are described by the *Adaptive Exponential Integrate-And-Fire Model* (Adex) (99). In this model, each neuron *i* is described by its membrane potential *V_i_*, which evolves according to the following equations:

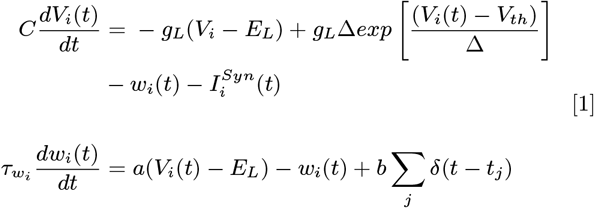

where *C* is the membrane capacitance, *g_L_* is the leakage conductance, *E_L_* is the leaky membrane potential, *V_th_* is the effective threshold, Δ is the threshold slope factor and 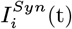 is postsynaptic current received by the neuron *i* (see next section). The adaptation current, described by the variable *w_i_*, increases by an amount *b* every time the neuron *i* emits a spike at times *t_j_* and decays exponentially with time scale *τ_w_* The subthreshold adaptation is governed by the parameter *a*.

During the simulations, the equation characterizing the membrane potential *V_i_* is numerically integrated until a spike is generated. Formally this happens when *V_i_* grows rapidly toward infinity. In practice, the spiking time is defined as the moment in which *V_i_* reaches a certain threshold (*V_th_*). When *V_i_* = *V_th_* the membrane potential is reset to *V_rest_*, which is kept constant until the end of the refractory period *T_ref_*. After the refractory period the equations start being integrated again.

In the developed network two types of cells were used: Regular Spiking (RS) excitatory cells and Fast Spiking (FS) inhibitory cells. The cell specific parameters are indicated in Table 1.

**Table 1.** Specific Neuron Model Parameters

| Parameter | RS | FS |
| --- | --- | --- |
| $V_{th}$ | -40 mV | -47.5 mV |
| $\Delta$ | 2 mV | 0.5 mV |
| $T_{ref}$ | 5 ms | 5 ms |
| $\tau_w$ | 500 ms | 500 ms |
| $a$ | 4 nS | 0 nS |
| $b$ | 20 pA | 0 pA |
| $C$ | 150 pF | 150 pF |
| $g_L$ | 10 nS | 10 nS |
| $E_L$ | -65 mV | -65 mV |
| $E_E$ | 0 mV | 0 mV |
| $E_I$ | -80 mV | -80 mV |
| $V_{rest}$ | -65 mV | -65 mV |

### Synaptic Models

The post-synaptic current received by each neuron *i* is composed by three components: two excitatory, referent to *AMPA* and *NMDA* synaptic channels, and one inhibitory, referent to *GABA_A_* channels.

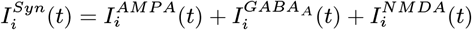

in which

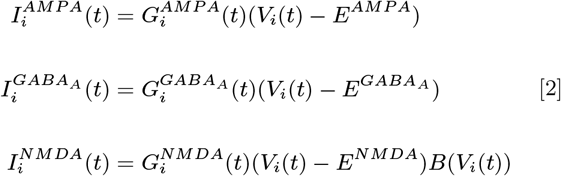

*E^AMPA^*= 0 mV, 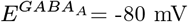 and *E^NMDA^*= 0 mV are the reversal potentials of *AM PA*, *GABA_A_* and *NMDA* channels. While the *AM PA* and *GABA_A_*-mediated currents are fast, the NMDA-mediated are considerably slower and present a complex relation with respect to the membrane potential (100–103). This complex relation, due to magnesium block, is accurately modeled by the phenomenological expression B(V) (104):

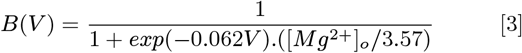

where [*Mg*^2+^]*_o_*= 1 mM is the external magnesium concentration (1 to 2 mM in physiological conditions).

Because of the fast dynamicas of *AMPA* and *GABA_A_* channels, their synaptic conductances (*G^X^* with X=*AMPA*, *GABA_A_*) are usually modeled to increase discontinuously by a discrete amount *Q^X^*, every time a presynaptic neuron spikes at time *t_k_*, and to subsequently decay exponentially with a decay time constant 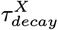 according to the following equation:

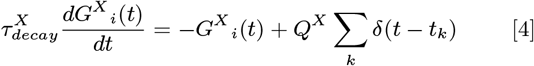

In which, Σ_*k*_ runs over all the presynaptic spike times. The synaptic time constantes used for *AMPA* and *GABA_A_* synapses are 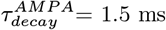 and 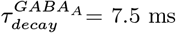.

NMDA channels synaptic conductances, *G^NMDA^*, because of their slow dynamics, are usually modeled as a bi-exponential function characterized by a rise time constant, 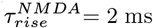, and a decay time constant 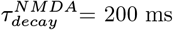, according to the following equation:

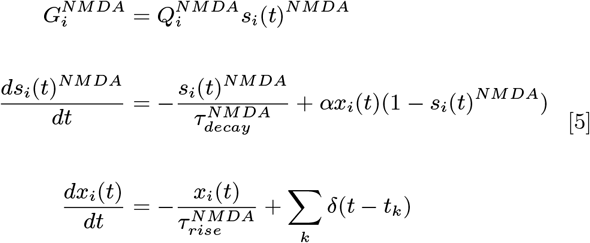

In which, 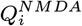 is the synaptic strength of the NMDA synapse towards the neuron *i*, *α*= 0.5/ms and *x*(*t*) is an auxiliary variable. The Σ_*k*_ runs over all the presynaptic spike times. Both, *s*(*t*)^*NMDA*^ and *x*(*t*), are adimensional.

Synaptic weights were chosen according to the parameter search expressed in Fig. 5. The the synaptic parameters of *AMPA* and *GABA_A_* synapses were chosen according to previous works (40, 105) (*Q^AMPA^*= 5 nS and 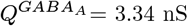). All synapses (*AMPA*, *GABA_A_* and *NMDA*) were delayed by time of 1.5 ms.

### Network Structure

The network developed in this work is composed of 5000 neurons (4000 RS and 1000 FS). Each neuron (RS or FS) was connected randomly to every other neurons in the network with a probability of 10%, receiving on average 500 excitatory synapses (mediated by both *AMPA* and *NMDA* channels) and 100 inhibitory synapses (mediated by *GABA_A_* channels).

### External Input

In addition to recurrent connections, each neuron received an external drive to keep the network active. This external drive consisted of *N_ext_* = 5000 independent and identically distributed excitatory Poissonian spike trains with a spiking frequency *μ_ext_*. These spike trains were sent to the network with a 10% probability of connection and were computed inside of the synaptic current term *I^AMPA^*, with a synaptic strength of 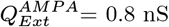. For gamma activity, the network was stimulated with a drive with *μ_ext_*= 3 Hz. For Asynchronous and Irregular activity, the network was stimulated with a drive with *μ_ext_*= 2 Hz. The external drive mimicked cortical input, like if the network was embedded in a much bigger one.

To test network responsiveness, an additional external input was included in the simulations. This external input, similar to the external drive, also consisted of *N_ext_* = 5000 independent and identically distributed excitatory Poissonian spike trains, connected to the network with a 10% probability. The difference of this input was its firing rate time dependence (*μ_ext_*(*t*)). The spiking frequency of the spike trains varied in a Gaussian manner, with a standard deviation of 50 ms and variable amplitude. These spike trains were computed inside of both synaptic current terms *I^AMPA^* and *I^NMDA^*, with a synaptic strengths of 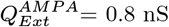, and 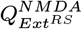 and 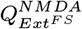 as indicated in each case.

### Block of NMDA channels: effect of NMDAR antagonists

In this work, we mimic the effect of NMDAR antagonists by changing the value of the NMDA synaptic weights *Q^NMDA^*. In Fig. 5 a possible trajectory in the parameter space generated by the action of NMDAR antagonists is depicted. This is the same trajectory indicated in Fig. 1A.

### Numerical simulations

All neural networks were constructed using Brian2 simulator (106). All equations were numerically integrated using Euler Methods and dt=0.1 ms as integration time step. The codes for each one of the three developed networks are available at ModelDB platform.

### Population activity: LFP model

To measure the global behavior of the neuronal population, we used a simulated Local Field Potential (LFP). This LFP was generated by the network, by means of a recent method developed by (107). This approach calculates the LFP by convolving the spike trains of the network with a Kernel that have been previously estimated from unitary LFPs (the LFP generated by a single axon, *uLFP*) measured experimentally. Since this method assumes a spatial neuronal displacement, to be able to apply it to our simulations, we randomly displaced part of the network (50 neurons) in 2-D grid, assuming that the electrode was displaced on its center and was measuring the LFP in the same layer as neuronal soma. The program code of the kernel method is available in ModelDB (http://modeldb.yale.edu/266508), using python 3 or the *hoc* language of NEURON.

### Power Spectrum

The Power Spectrum of the simulated LFP was calculated by means of the Welch’s method, using a Hamming window of length 0.25 seconds and 125 overlapping points. We used the Python-based ecosystem Scipy function *signal.welch* to do our calculations.

### Synaptic Charge

The synaptic charge (AMPA or NMDA) of each neuron is defined as the area under the curve of the average synaptic current (shaded areas of Fig. 6A or B), which was calculated from the presynaptic input time until 10 ms after it.

### Responsiveness

The level of *responsiveness* (*R*) of a network, due to a stimulus (*S*) in a time window of duration *T*, is defined as the difference between the total number of spikes generated by the whole network due to an stimulus 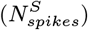 and the total number of spikes generated in the absence of the stimulus (*N_spikes_*), normalized by the network size (total number of neurons *N_n_*) and the duration of the time window *T*.

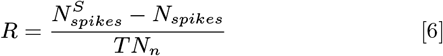

## Acknowledgments

This research was supported by the Centre National de la Recherche Scientifique (CNRS) and the European Community (Human Brain Project, H2020-785907, H2020-945539). E.S. acknowledges a PhD fellowship from the École des Neurosciences de Paris (ENP) and from the Fondation pour la Recherche Médicale (FRM, grant FDT202012010566) and a financial support from La Fondation des Treilles.

